# GenomeDelta: detecting recent transposable element invasions without repeat library

**DOI:** 10.1101/2024.06.28.601149

**Authors:** Riccardo Pianezza, Anna Haider, Robert Kofler

## Abstract

To evade repression by the host defense, transposable elements (TEs) are occasionally horizontally transferred (HT) to naive species. TE invasions triggered by HT may be much more abundant than previously thought. For example, previous studies in *Drosophila melanogaster* found 11 TE invasions over 200 the past years. A major limitation of current approaches for detecting recent invasions is the necessity for a repeat-library, which is notoriously difficult to generate. To address this, we developed GenomeDelta, a novel approach for identifying sample-specific sequences, such as recently invading TEs, without prior knowledge of the sequence. It can thus be used with model and non-model organisms. As input, GenomeDelta requires a long-read assembly and short-read data. It will find sequences in the assembly that are not represented in the short read data. Beyond identifying recent TE invasions, GenomeDelta can detect sequences with spatially heterogeneous distributions, recent insertions of viral elements and recent lateral gene transfers. We thoroughly validated GenomeDelta with simulated and real data from extant and historical specimens. Finally, we demonstrate that GenomeDelta can reveal novel biological insights: we discovered the three most recent TE invasions in *Drosophila melanogaster* and a novel TE with a geographically heterogeneous distribution in *Zymoseptoria tritici*.

## 1 Introduction

Transposable elements (TEs) are short DNA sequences capable of increasing their copy numbers within a host genome. They are common in many organisms and often make up a large part of their genome [Chénais et al., 2012, Wicker et al., 2007]. While some TEs may confer benefits to hosts [Eickbush and Eickbush, 1995, Nelson et al., 2023], the majority of TE insertions are likely deleterious [Elena et al., 1998, Pasyukova et al., 2004].

Consequently, host genomes have evolved elaborate defense mechanisms, frequently involving small RNAs [Sarkies et al., 2015]. TEs can evade host silencing through horizontal transfer (HT), i.e. the transmission to naive species that lack the TE [Peccoud et al., 2017, Scarpa et al., 2023, Kofler et al., 2015, Signor et al., 2023]. HT is a common phenomenon in prokaryotes [Koonin et al., 2001], but recent studies suggest that HT (especially of TEs) is also prevalent in eukaryotic organisms [Peccoud et al., 2017, Zhang et al., 2020].

The evidence for HT of TEs has typically been indirect. Such evidence includes a patchy distribution of the TE among closely related species or a high similarity between the TE of the donor and recipient species, which is frequently quantified by the synonymous divergence of the TE Peccoud et al. [2018], Wallau et al. [2012].

However, for TEs that have spread very recently, direct evidence of the recent invasion can be obtained, for example when the sequence of the TE is absent in older samples but present in more recently collected ones.

To illustrate, the *P-element* invaded *D. melanogaster* populations between 1950 and 1980 [Kidwell, 1983, Anxolabéhère et al., 1988]. Consequently, sequences with similarity to the *P-element* are absent in natural *D. melanogaster* strains collected before 1950 but present in strains collected after 1980. It is possible that such recent invasions are common. In *D. melanogaster*, an important model organism for studying TE dynamics, 11 different TE families invaded natural populations during the last 200 years [Kidwell, 1983, Anxolabéhère et al., 1988, Bucheton et al., 1992, Bonnivard et al., 2000, Schwarz et al., 2021, Pianezza et al., 2023, Scarpa et al., 2023, Pianezza et al., 2024]. It is feasible that other organisms might also experience a high rate of recent invasions.

To obtain direct evidence for recent TE invasion, it is necessary to compare the sequencing data from old and recent samples. Recent samples can be collected from natural populations, whereas old samples may be derived from several sources, including old laboratory strains, genomes of historical specimens from museums or ancient DNA extracted from archaeological remains [Raxworthy and Smith, 2021, Orlando et al., 2021, Schwarz et al., 2021]. The number of species with sequencing data from historical specimens is rapidly increasing [Benham and Bowie, 2023] (for example *Anopheles* sp. [Korlević et al., 2021], *Apis mellifera* [Parejo et al., 2020], *Columba livia* [Hernández-Alonso et al., 2023] and *Canis lupus* [Bergström et al., 2022]). Therefore, it is in principle feasible to discover direct evidence for recent TE invasions in an increasing number of species. However, discovering recent invasions with existing approaches typically requires comparing the copy numbers of known TE families in old and young samples [e.g. Scarpa et al., 2023, Schwarz et al., 2021]. These approaches thus require prior knowledge of the sequences of the TEs, i.e. a repeat library. Generating repeat libraries is notoriously difficult, requiring extensive manual curation [Hoen et al., 2015, Rodriguez and Arkhipova, 2022, Goubert et al., 2022]. This issue is further compounded by the fact that even for the few species for which a high-quality repeat library is available, the library may be incomplete and not contain the sequences of TEs that have spread very recently. For example, the high quality repeat library of *D. melanogaster* [Quesneville et al., 2005] lacks the sequence of the retrotransposon *Spoink*, which spread in natural populations between 1983 and 1993. This is because the reference strain used for generating the repeat library was likely collected prior to that period. This is part of the reasons why the *Spoink* invasion was only recently discovered, several years after the invasion [Pianezza et al., 2023].

The development of an approach that enables identification of recent TE invasions independent of repeat libraries would represent a substantial conceptual advance in the field. For this reason we, developed GenomeDelta. GenomeDelta is based on the idea that recent invasions will lead to sequences that are present in recently collected samples (i.e. after the invasion) but absent in old samples (i.e. before the invasion). As input, GenomeDelta requires a high quality assembly of the recently collected sample and short-read data of the old sample. GenomeDelta then identifies sequences that are present in the assembly but absent in the short read data. As this approach does not require prior knowledge about the sequences, it allows to comprehensively identify sample-specific sequences (e.g. TEs that invaded recently) in model and non-model organism. Apart from finding recent TE invasions, GenomeDelta may also be used to detect sequences showing a geographically heterogeneous distribution, such as the TE *Styx* in *Z. tritici* [Feurtey et al., 2023b], recent endogenous retrovirus insertions [Gilbert and Belliardo, 2022] and recent lateral gene transfer [Dunning et al., 2019]. We thoroughly validated our novel tool with simulated and real data. We also provide a detailed manual and a walkthrough. Finally, we show that GenomeDelta can be used to gain novel biological insights. With GenomeDelta, we discovered three novel TE invasion *D. melanogaster* in the last three decades and a novel TE (*Rosetta*) with a spatially heterogeneous distribution in *Z. tritici*.

## 2 Results

### 2.1 GenomeDelta

We developed GenomeDelta (GD) to identify genomic sequences that are present in one sample (*P* presence) and absent in another sample (*A* absence). Given the two sets of genomic sequences ‘*P* ‘ and ‘*A*’, GenomeDelta aims to identify the set of sample-specific sequences *P* − *A*. Note that GenomeDelta does not compute the set *A* − *P*.

As input, GenomeDelta requires a high-quality assembly for sample *P* and short read data for sample *A*. GenomeDelta is based on the idea that sequences that are present in *P* and absent in *A* (i.e. *P* − *A*) can be identified as coverage gaps when short reads of *A* are aligned to an assembly of *P* (fig. 1A).

**Figure 1:**
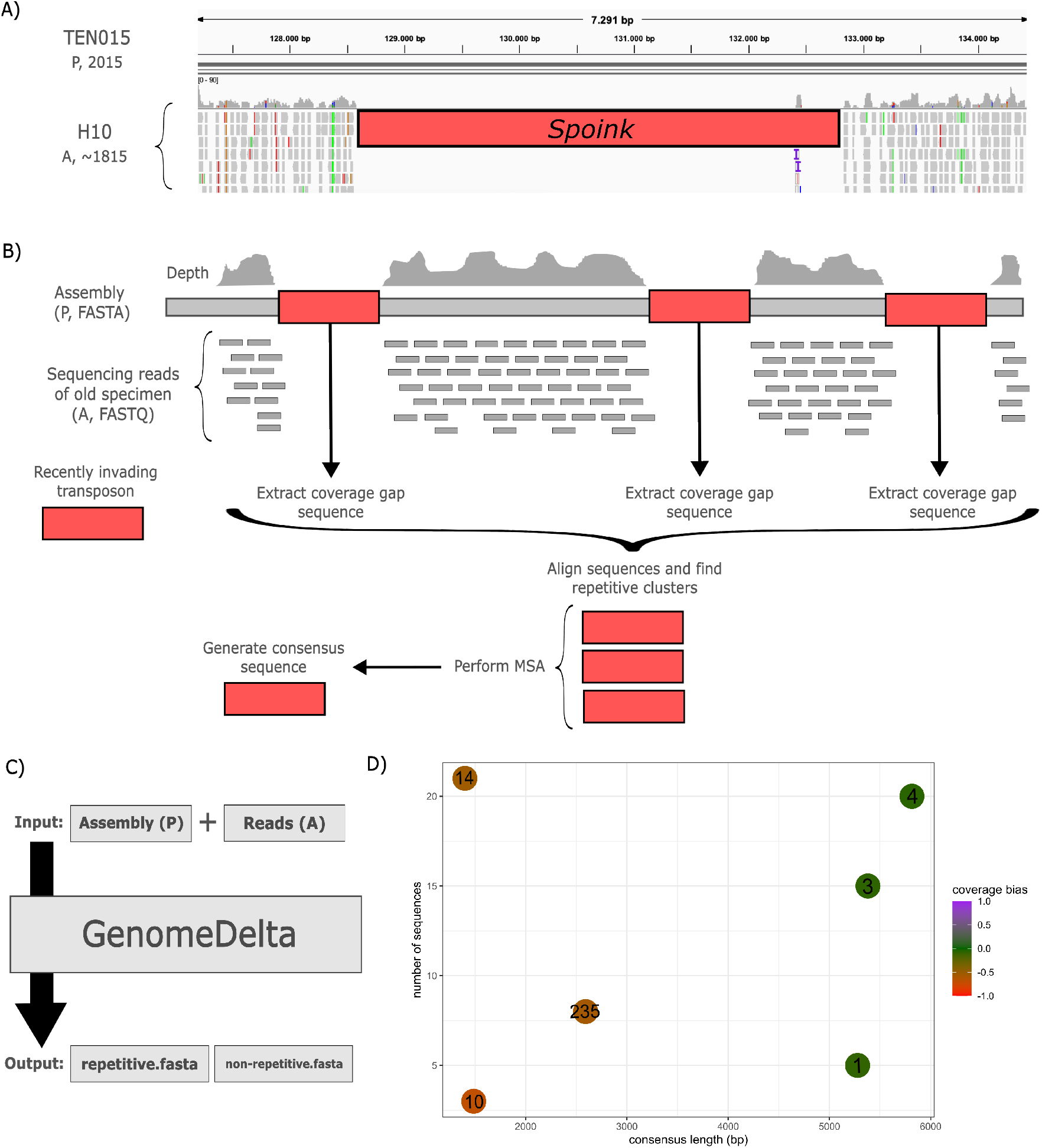
Overview of GenomeDelta (A) Recent TE invasions will lead to coverage gaps when reads of a sample collected before the invasion (H10, collected in 1815) are aligned to the assembly of a sample collected after the invasion (TEN015, collected in 2015). The coverage gap in this example is due to the retrotransposon *Spoink*, which invaded *D. melanogaster* populations between 1983-1993 [Pianezza et al., 2023]. (B) Workflow of GenomeDelta. Reads (FASTQ) of a sample *A* (absence) are aligned to an assembly (FASTA) of another sample *P* (presence). The sequences of coverage gaps are extracted, similar sequences are clustered, a multiple sequence alignment is performed and consensus sequences are generated. Finally, the sequences that are present in *P* but absent in *A* are reported. (C) Overview of the input and output of GenomeDelta. As output, two fasta-files are generated, one with the consensus sequences of repetitive elements and one with the sequences of non-repetitive elements. (D) Graphical output generated by GenomeDelta, providing an intuitive overview of the sample-specific repetitive sequences. The length, the copy number and the coverage bias (see text) is shown for each identified repetitive sequence.

One major field of application for GenomeDelta is the identification of novel TE invasions. TE invasions add novel sequences to the genome that are present in samples collected after the invasion (*P*) but absent in samples collected before the invasion (*A*). Another major use-case of GenomeDelta is the identification of sequences (TEs) that are present in one geographic region (*P*) but absent in another (*A*). For example *KoRV* (koala retrovirus) insertions are present in the genomes of koalas sampled from the North but not in all the koalas from the South of Australia [Tarlinton et al., 2006]. In summary, GenomeDelta may be used to identify sequences (repetitive or non-repetitive) showing a spatial or temporal heterogeneous presence/absence pattern.

To identify the sample-specific sequences (*P* − *A*), GenomeDelta aligns the sequencing reads of *A* to the assembly of *P* and computes the coverage. Next, GenomeDelta identifies coverage gaps, extracts the sequences of the gaps, groups them by sequence similarity (e.g. the different insertions of a TE family), performs a multiple sequence alignment (MSA) and reports the consensus sequences (fig. 1B). Separate results are reported for repetitive and non-repetitive sample-specific sequences (fig. 1C). Finally, GenomeDelta estimates the reliability for each of these sample-specific sequences (*P* − *A*) by computing a coverage bias score. The coverage bias is estimated as 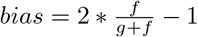 where *f* is the coverage in the regions flanking the coverage gap (10, 000 bp in each direction) and *g* the average genomic coverage. The bias ranges from −1 to 1, where 0 indicates an unbiased coverage and −1 and 1 a highly biased coverage (either highly decreased or increased; supplementary fig. S1). As output, the sample specific-sequences are provided as two fasta-files, one for the consensus sequences of repetitive elements and one for the non-repetitive sequences. Additionally, a bed file with the genomic coordinates and the coverage bias of each coverage gap is reported. To provide an intuitive graphical overview, GenomeDelta also generates a summary plot, showing for each sample-specific repetitive sequence the copy number, the length and the coverage bias (fig. 1D).

GenomeDelta can be easily installed with conda ([Anaconda, 2012]). A detailed manual and a walkthrough with data from *D. melanogaster* are available. GenomeDelta is distributed under the Open Software License v.2.1.

### 2.2 Validation

We thoroughly validated GenomeDelta with simulated and real data. For validations with simulated data, we used a chromosome sequence of *D. melanogaster* (chromosome arm 2R) as template and inserted 25 copies of a randomly generated sequence with a length of 5000 bp into this template. We thus obtain an artificial sequence with (*P*) and without TEs (i.e. the template, *A*). Next, we generated artificial short-reads for the sequence without TEs (*A*). We simulated artificial reads yielding different coverages and coverage distributions (uniform coverage and random position of reads; table 2.2). We also used Gargammel [Renaud et al., 2017] to simulate reads mimicking properties of ancient DNA (i.e. fragmentation, cytosine deamination, bacterial contamination [Orlando et al., 2021]). Finally, we plugged the artificial reads into GenomeDelta to identify the sample-specific sequences. To evaluate the performance of GenomeDelta, we computed the true positive rate, the false positive rate, the length of the identified consensus sequence and the similarity between the observed consensus sequence and the simulated sequence (table 2.2). The 25 artificial insertions were detected in all scenarios. The obtained consensus sequence was 100% identical to the simulated sequence, and the length of the consensus sequence was close to the simulated 5000bp. With a low coverage, especially when properties of ancient DNA were simulated, the false positive rate was elevated and the length of the consensus sequence deviated from the expected one (i.e. 5000; table 2.2). However, with a coverage of ≥ 5, no false positives were found and the observed length was close to the expected one (table 2.2). To test the performance of GenomeDelta with short sequences, we performed an additional validation with a sequence of length 1000bp. With such a short sequence, GenomeDelta identified more false positive sequences when the coverage was low (i.e 1x, supplementary table S1). Finally, we simulated reads with different lengths and sequencing error rates. The error rate of the reads and the read length had little impact on the performance of GenomeDelta (supplementary tables S2, S3).

Our validations with simulated data suggest that GenomeDelta accurately identifies sample specific-sequences with a length *>* 1000bp. Furthermore, the short-read data (of sample *A*) ought to have a minimum coverage of 5.

**Table 1:** Validation of GenomeDelta with simulated data. We introduced 25 copies of an artificial TE with a size of 5000bp into a template sequence and tested if the TE sequence was accurately identified with GenomeDelta. As input, we used the template with the TEs (*P*) and artificial reads simulated from the template without TEs (*A*). We simulated different read coverages using either a uniform or a heterogeneous coverage (random position of reads). We also simulated properties of ancient DNA with Gargammel. We evaluated the number of true positive insertions (*TP*), the number of false positive insertions (*FP*_*r*_, *FP*_*nr*_: repetitive or non-repetitive), the length of the reported consensus sequence [len. (bp)] as well as the sequence similarity between the reported consensus sequence and the simulated insertion [sim (%)].

| method | coverage | *TP* | *FP<sub>nr</sub>* | *FP<sub>r</sub>* | len. (bp) | sim (%) |
| --- | --- | --- | --- | --- | --- | --- |
| uniform | 1 | 25 | 8 | 0 | 5,000 | 100 |
| uniform | 5 | 25 | 0 | 0 | 4,998 | 100 |
| uniform | 10 | 25 | 0 | 0 | 4,998 | 100 |
| random | 1 | 25 | 607 | 2 | 5,251 | 100 |
| random | 5 | 25 | 0 | 0 | 4,998 | 100 |
| random | 10 | 25 | 0 | 0 | 4,997 | 100 |
| gargammel | 1 | 25 | 2,410 | 17 | 5,435 | 100 |
| gargammel | 5 | 25 | 0 | 0 | 5,000 | 100 |
| gargammel | 10 | 25 | 0 | 0 | 4,997 | 100 |

Next, we validated GenomeDelta with real genomic data from an insect (the fruit fly, *D. melanogaster*) and a mammal (the koala, *Phascolarctos cinereus*).

Based on various approaches, ranging from phenotyping (hybrid dysgenesis) to whole genome sequencing, previous works showed that seven TEs spread in *D. melanogaster* populations between 1800 and 1980 [Anxolabéh`ere et al., 1988, Daniels et al., 1990, Bonnivard et al., 2000, Bucheton et al., 1992, Kidwell, 1983, Schwarz et al., 2021, Scarpa et al., 2023]. These works showed that *Blood, 412, Opus* spread between 1850 and 1930, *Tirant* around 1935 followed by the *I* -element, *Hobo*, and the *P* -element (fig. 2A). We tested whether GenomeDelta identifies these known invasions when short-read data of a strain collected around 1815 (H10) are aligned to the assembly of a strain collected around 1975 (Pi2) [Shpak et al., 2023, Wierzbicki et al., 2022]. Note that the short-read data are derived from a strain kept for ≈200 years in a museum, thus the sequencing reads are fragmented with an average size of around 50bp [Shpak et al., 2023]. GenomeDelta detected 27 repetitive sequences that are specific to Pi2 (fig. 2B). A blast search of the consensus sequences against a TE library [Chakraborty et al., 2021, Quesneville et al., 2005] revealed that the 7 TE that invaded *D. melanogaster* recently are the most notable outliers in terms of length, copy numbers and coverage bias (fig. 2B). Note that several fragments are reported for some TEs (*Blood, I* -element, *Tirant* ; fig. 2B). This can be explained by the fact that degraded fragments of these TEs, likely the remnants of ancient invasions, are present in all genomes, including the genome of the strain sampled at 1815 [Schwarz et al., 2021, Scarpa et al., 2023]. Reads derived from these fragments may be mapped on the new TE insertions, thus fragmenting the coverage gap generated by the new insertion. The other sample-specific sequences were less promising, as for example seen by the high coverage bias. Closer inspection revealed that many of these sequences were located in telomeric regions where few reads aligned in Pi2 (supplementary table S4). GenomeDelta thus identified the 7 TEs that invaded *D. melanogaster* between 1815 and 1975 based on sequencing reads from historical specimens [Shpak et al., 2023].

**Figure 2:**
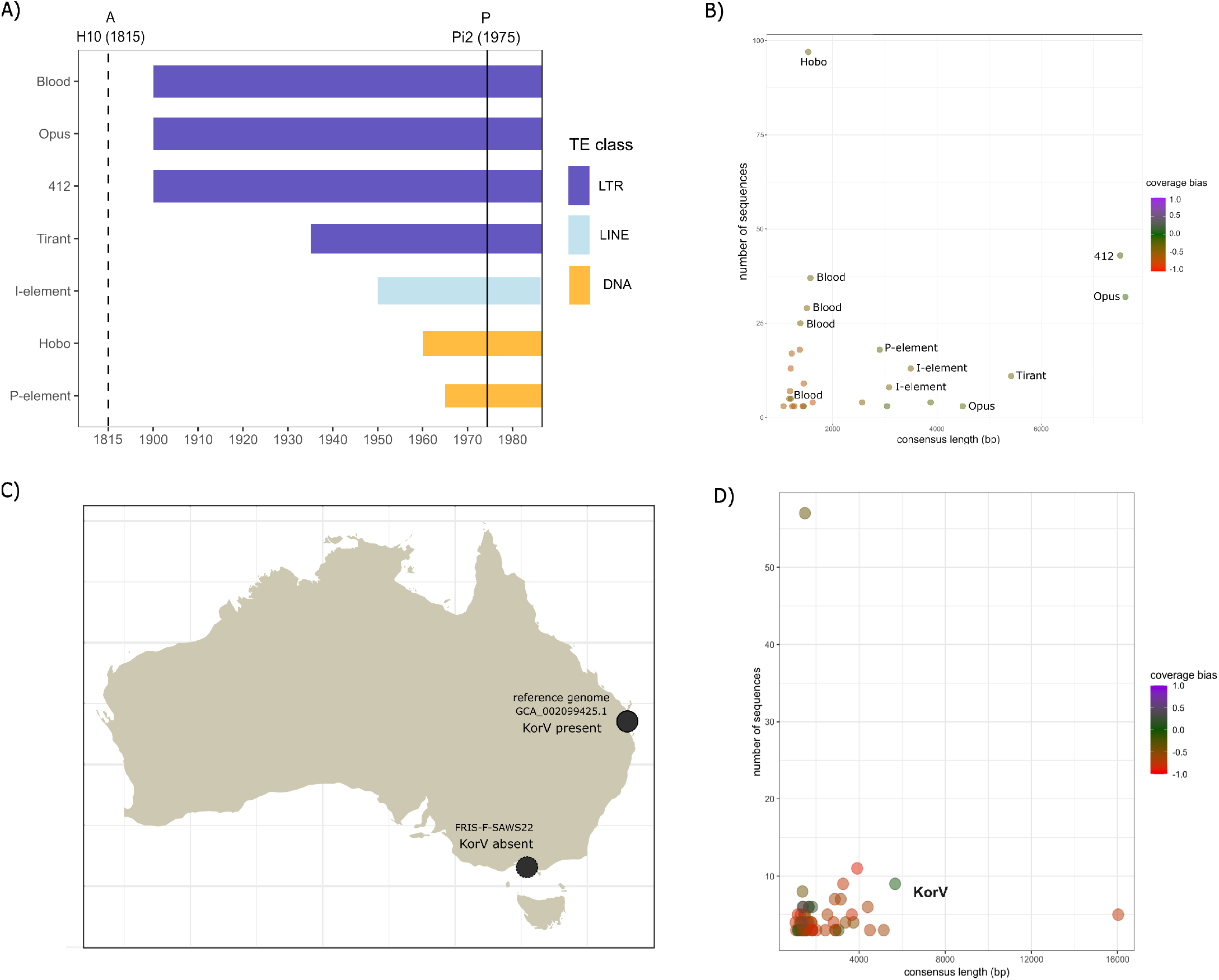
Validation of GenomeDelta with real data A) Overview of the invasion history of *D. melanogaster* until 1975 as revealed by previous works. B) Overview of the sequences identified by GenomeDelta when a sample collected around 1815 (H10) is compared to a strain collected in 1975 (Pi2). Note that the identified sequences correspond to the recent invaders. C) Previous work shows that KoRV insertions are present in the genomes of koalas sampled in the North but not in the South of Australia. D) Overview of sequences identified by GenomeDelta when short reads from a southern koala (FRIS-M-SAWS21) are aligned to an assembly of a northern koala (GCA 002099425.1). Note that the best candidate sequence (i.e. coverage bias close to zero) corresponds to KoRV.

We next evaluated the influence of the coverage on the performance of GenomeDelta with real data. The sample used to identify the 7 invasions, H10, yielded a coverage of 33x [Shpak et al., 2023]. We subsampled the number of reads of H10 to coverages of 10x, 5x and 1x, and again used GenomeDelta to identify sequences specific to Pi2 (supplementary fig. S2). With a coverage of 10x, GenomeDelta identified 5 out of the 7 TEs (*Blood* and *Tirant* were missing). With a coverage of 5x, we identified the same 5 out of the 7 TE, but additionally *Hobo* and *Opus* were fragmented into two clusters. With a coverage of 1x, just one short fragment of the *I-element* was found (supplementary fig. S2). In agreement with the validation with artificial data, the subsampling of real data (from historical specimens) suggest that a coverage *>* 5 should be used with GenomeDelta.

We next tested the performance of GenomeDelta with a mammal, i.e. the koala (*Phascolarctos cinereus*). Insertions of the koala retrovirus (KoRV) have been found in the genomes of koalas sampled from the North of Australia but not in koalas samples from the South [Tarlinton et al., 2006]. KoRV may be at the transition stage between an exogenous element (e.g. virus) and a vertically transmitted endogenous element (i.e. a transposable element [Tarlinton et al., 2006]). We tested whether GenomeDelta manages to identify KoRV, by comparing short-read data from a southern koala (FRIS-M-SAWS21 [Hogg et al., 2023]) to the assembly of a northern koala (i.e. the reference genome GCA 002099425.1, which is based on a koala from the Sunshine coast; fig. 2C). GenomeDelta identified several sample-specific repetitive sequences (fig. 2D). The most promising sequence (low coverage bias, high copy number and substantial length) corresponds to KoRV (fig. 2D). Compared to *D. melanogaster*, we find more false positive sequences (57 in koalas and 11 in *D. melanogaster* ; fig. 2B,D). This can likely be explained by the much larger genome size of koalas as compared to *D. melanogaster* (3500 Mb in koalas and 200 Mb in *D. melanogaster* [Johnson et al., 2018, Celniker and Rubin, 2003]). As a control, we also compared two koalas from a northern population with GenomeDelta (Sunshine-Coast-M-79817 and the reference genome) and did not find any sequences matching with KoRV (supplementary fig. S3). In agreement with previous work, GenomeDelta thus identified the integration of KoRV in the genomes of northern koalas but not in southern koalas [Tarlinton et al., 2006].

In summary we thoroughly validated GenomeDelta with artificial and real data. We also showed that GenomeDelta may be used with historical DNA and that a minimum coverage of 5 should be targeted.

### 2.3 Novel biological insights

Finally, we provide two examples illustrating how GenomeDelta can be used to generate novel biological insights. First, we identified three novel TE invasions in *D. melanogaster* with GenomeDelta (fig. 2A; [Pianezza et al., 2024]). We used GenomeDelta to align reads from a sample collected in the early 1800s (H10) to the assembly of a strain collected in 2016 (TOM007). As expected, we found all the previously described TE invasions that occurred between 1810 and 1975 (fig. 2A, B) and *Spoink*, which spread between 1983 and 1993 [Pianezza et al., 2023]. Surprisingly, GenomeDelta also discovered three novel TE invasions in *D. melanogaster* : *MLE, Souslik* and *Transib1* (fig. 3A). We are describing these novel invasions in detail in a separate work [Pianezza et al., 2024]. Here, we use these three novel invasions to showcase how candidates identified with GenomeDelta may be further validated and investigated. We will thus start with the raw results provided by GenomeDelta. As a first step, we investigated the coverage bias. Since the three novel sequences have a low bias (i.e. close to zero), they may be considered promising candidates (fig. 3A blue). Next, a blast search revealed that these sequences correspond to three different annotated TEs, a “Micropia-like” element (*MLE*) described in *D. melanogaster* (GenBank: MN418888.1), *Transib1* described in *D. melanogaster* and *D. simulans*, and to *Souslik* identified in *D. simulans* (GenBank: BK008880.1). For some TEs, such as *Blood*, the consensus sequence identified by GenomeDelta may be fragmented or incomplete (fig. 3) To obtain the complete consensus sequence of the novel TEs, we extracted the sequence of each insertion together with the flanking sequences (3000bp) using bedtools. Next, we performed a multiple sequence alignment of these sequences (TE + flanking region) with MUSCLE and constructed a novel consensus sequence using the GenomeDelta script ‘MSA2consensus.py’, which employs a simple majority rule for generating consensus sequences. While the consensus sequences of *MLE* and *Souslik* remained largely unchanged, the length of the consensus sequence of *Transib1* increased from 1323 to 3030 bp. This shows that the initial consensus sequence of *Transib1* extracted by GenomeDelta was incomplete. This finding can be attributed to the presence of degraded fragments of *Transib1*, likely remnants of past invasions, in all *D. melanogaster* strains ([Kapitonov and Jurka, 2003, Pianezza et al., 2024]). Reads from ancient *Transib1* fragments may be misaligned to the recent *Transib1* insertions, thus interfering with GenomeDelta’s identification of coverage gaps. Next, we identified the LTR sequence of *MLE* and *Souslik* and the inverted repeat sequences of *Transib1* using BLAST. Based on long-read assemblies, we confirmed the presence of these three TEs in strains collected around 2015 and their absence in strains collected before 1975 [Pianezza et al., 2024]. We inferred the exact timing of these three invasions using 585 *D. melanogaster* samples collected during the last 200 years [Pianezza et al., 2024]. Finally, we aimed to identify the species that acted as donor of the horizontal transfer triggering the invasions, by analysing the genomes of 1400 arthropod species[Pianezza et al., 2024]. By identifying three recent TE invasions in *D. melanogaster*, we demonstrated that GenomeDelta can be used to generate novel insights about sequences showing a temporally heterogeneous distribution [Pianezza et al., 2024].

**Figure 3:**
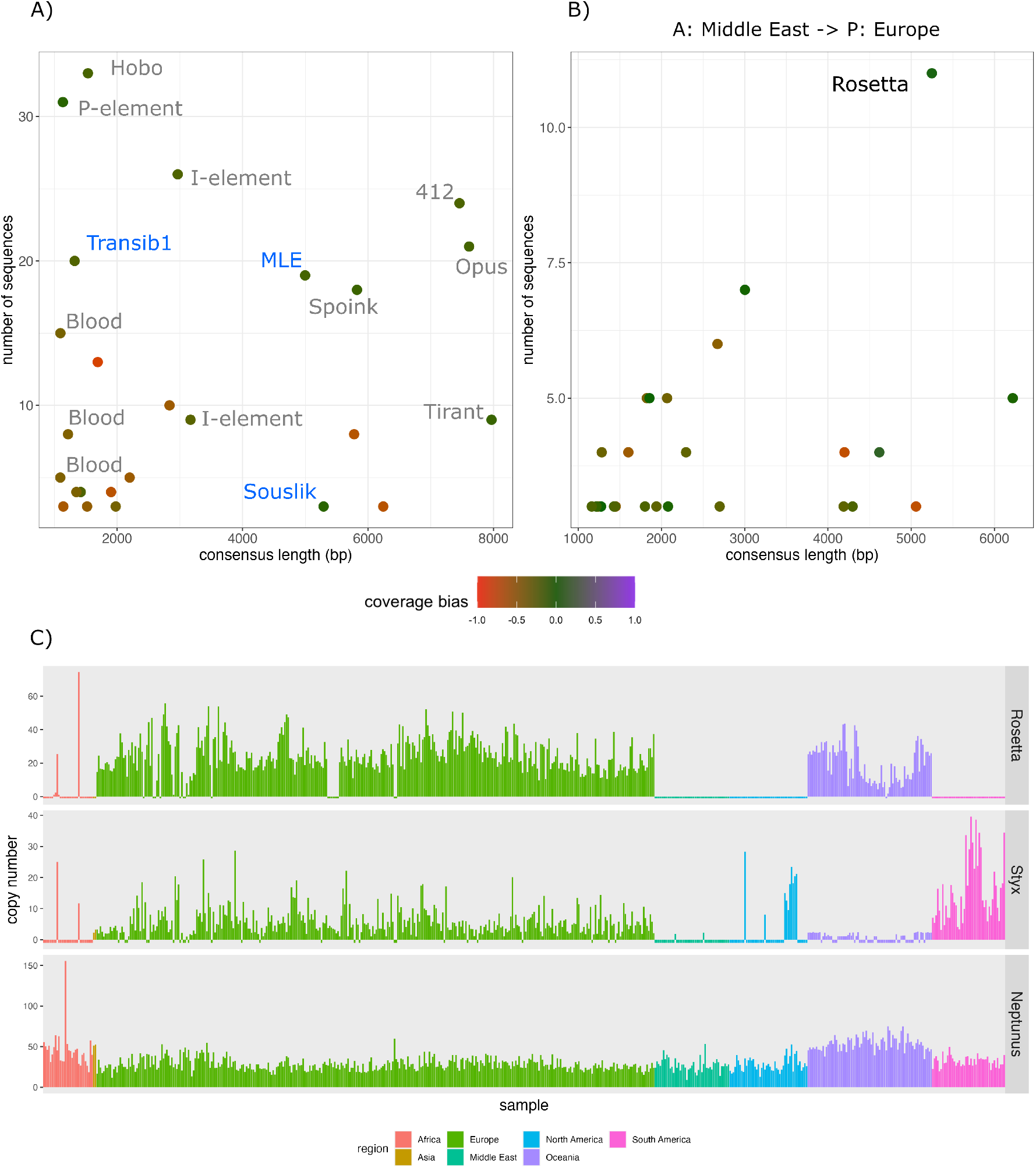
GenomeDelta can be used to generate novel biological insights (a) GenomeDelta identifies three novel TE invasions in *D. melanogaster* (blue). Short-read data of a strain collected in the early 1800s (*A*: H10) were aligned to the assembly of a strain collected in 2016 (*P* : TOM007). Previously identified TE invasions are in black. (b) GenomeDelta identifies a novel TE (*Rosetta*) with a spatially heterogeneous distribution in *Z. tritici*. Short reads data of a sample from Iran (*A*: SRR5194593) were aligned to the assembly of a sample from the Netherlands (*P* : GCA000219625.1). (c) Copy number of *Rosetta* in *Z. tritici* samples collected from different geographic regions. As controls, a TE with a known spatially heterogeneous distribution (*Styx*) and a TE present in all strains (*Neptunus*) are shown. TE copy numbers were estimated with DeviaTE.

We next tested whether GenomeDelta can also be used to generate novel insights about sequences showing a spatially heterogeneous distribution. We utilized the publicly available genomic resources for the important crop pathogen *Zymnoseptoria tritici* (see files S2and S1), which causes septoria leaf blotch, one of the most important diseases of wheat [Stukenbrock et al., 2010]. *Z. tritici* is native to the Middle East, but spread to all continents between 500 and 200 years ago [Feurtey et al., 2023a]. Elegant recent work revealed that several TEs (*Styx, Juno, Deimos* and *Fatima*) show a spatially heterogeneous distribution in *Z. tritici*, where for example the TE *Styx* is present in populations from Europe but not in the Middle East [Feurtey et al., 2023a] (see also fig. 3C). The authors attributed this heterogeneous distribution of the TEs to spatial differences in the efficiency of the genomic host defenses against TEs (repeat-induced point mutations [Van Wyk et al., 2021]). Using GenomeDelta, we aligned short reads from a Middle East sample (SRR5194593) to the assemblies of 15 *Z. tritici* strains collected from diverse continents. As expected, a comparison between the sample from the Middle East and Australia identified TEs that were previously shown to have a geographic heterogeneous distribution (*Styx, Juno, Deimos* and *Fatima*, supplementary fig. S4; [Feurtey et al., 2023a]). Interestingly, by comparing a sample from the Middle East to a sample from the Netherlands, GenomeDelta revealed a novel TE with a heterogeneous distribution, i.e. *Rosetta* (fig. 3B). Although *Rosetta* has been previously annotated in *Z. tritici*, a spatially heterogeneous composition has not been described before [Badet et al., 2020, Feurtey et al., 2023a]. To further investigate the abundance of *Rosetta* in samples from different geographic regions, we used DeviaTE [Weilguny and Kofler, 2019]. DeviaTE estimates the copy numbers of TEs by normalizing the coverage of TEs to the coverage of single copy genes. We found that *Rosetta* is most prevalent in Oceania and Europe but largely absent in Africa, the Middle East, North and South America (fig. 3C). As control, we also analysed the abundance of *Styx* with DeviaTE. In agreement with previous work, *Styx* was found in Europe and the Americas but not in Africa and Oceania (fig. 3C). As a further control we included a TE (*Neptunus*) present in all populations. A high number of *Neptunus* insertions can be found in all analysed samples (fig. 3C). GenomeDelta thus identified *Rosetta* as a novel TE with a spatially heterogeneous distribution in *Z. tritici*. Since the distribution of *Styx* and *Rosetta* varies among regions (e.g. *Rosetta* but not *Styx* is present in Oceania), it is questionable whether differences in the efficiency of the genomic defense as proposed previously [Feurtey et al., 2023a] can account for the spatially heterogeneous distribution of the TEs. Differences in the host defence ought to affect all TEs equally. An alternative hypothesis might be that *Styx* and *Rosetta* recently spread in *Z. tritici* following a horizontal transfer. The differences in distribution of *Styx* and *Rosetta* can then simply be explained by different geographic origins of the horizontal transfer that triggered the invasions or by differences in invasion routes caused by stochastic migration events transmitting the TE between the populations.

In summary, we showed that GenomeDelta can be used to gain novel biological insight into spatially (*Z. tritici*) and temporally (*D. melanogaster*) heterogeneous distributions of TEs.

## 3 Discussion

In this work we presented GenomeDelta, a tool designed to detect genomic sequences that are present in one sample but absent in another one. We thoroughly validated GenomeDelta with both artificial and real data, and showed that GenomeDelta can be used to generate novel biological insights by detecting recent TE invasions in *D. melanogaster* and identifying a TE with a geographically heterogeneous distribution in *Z. tritici* (fig. 3).

As major advantage, GenomeDelta can be used to identify sample-specific sequences without prior knowledge about the sequences. Such sample specific sequences are of biological interest and could be generated due to varying processes, ranging from horizontal transfer, to differences in the host defence against foreign DNA and to locally restricted negative selection against some sequences. One important use-case for finding sample specific sequences is the identification of recent TE invasions, where a TE is present in recent but absent in older samples. Previous approaches for finding such recent TE invasions required a comprehensive repeat library, which enables estimating copy number differences of repeats among samples of interest. Creating such repeat libraries is notoriously difficult and requires substantial manual curation [Hoen et al., 2015, Rodriguez and Arkhipova, 2022, Goubert et al., 2022]. Even more problematic is that a single repeat library for a species may not be sufficient, as some TEs may not be present in the library. For example, several TEs that invaded *D. melanogaster* recently (*Spoink, MLE* and *Souslik*) are not present in the high quality repeat library of *D. melanogaster* (these TEs spread after the strain used for generating the repeat library was collected [Pianezza et al., 2024, Quesneville et al., 2005]). A comprehensive identification of all TE invasions would thus require a ‘pan repeat library’ for a species, i.e. a library comprising the repeat sequences of a large number of strains sampled at different times from diverse geographic regions. Generating up-to-date repeat-libraries is thus a major bottleneck with conventional approaches for finding recent TE invasions. The identification of sample-specific sequences with GenomeDelta, independent of a repeat library, is thus a major conceptual advance, that enables identifying all recent TE invasions in model and non-model organisms.

GenomeDelta identifies sample-specific sequences (*P* − *A*) by aligning short-reads (sample *A*) to an assembly (sample *P*). Our validations indicate that historical DNA can be used (sample *A*) and that the short-read data (sample *A*) should have a minimum coverage of 5. This coverage requirement of 5 may be a limitation for some projects where shallow sequencing was performed for several samples. In this case, one workaround may be to simply pool the reads of all samples with shared properties (e.g. to pool all samples from the same geographic region or the same decade).

Another limitation is that GenomeDelta solely identifies sequences with qualitative differences, i.e. being present in some samples and absent in others. It will not identify sequences having quantitative differences in copy numbers among samples.

Perhaps the major limitation of GenomeDelta is the requirement for a high quality genome assembly (sample *P*). It is therefore not possible to identify sequences specific to samples from which a high quality assembly cannot be generated, such as historical samples, which typically yield only highly fragmented DNA [Raxworthy and Smith, 2021]. However, the need for a high quality assembly is not unique to GenomeDelta; such assemblies are also indispensable for approaches that rely on a repeat library. In fact, without a highquality assembly, it is impossible to construct a comprehensive repeat library.

Despite these limitations, we anticipate that GenomeDelta may be used for a wide range of applications. Our primary motivation for designing GenomeDelta was to discover recent TE invasions. For example, we used GenomeDelta to discover the three most recent TE invasions in *D. melanogaster* (fig. 3A; [Pianezza et al., 2024]). As another application, GenomeDelta might be used to identify sequences occurring in populations of some regions but not in others. We demonstrate this by using GenomeDelta to discover a novel TE with a geographically heterogeneous composition in *Z. tritici* (*Rosetta*; fig. 3B,C). Such a geographically heterogeneous distribution of TEs might result from spatial differences in the efficiency of the genomic host defence against TEs [Feurtey et al., 2023a]. We propose different invasion routes of the TEs following a horizontal transfer might also account for the observed heterogeneity. Finally, GenomeDelta could also be used to identify sample-specific non-repetitive sequences such as recent endogenous retrovirus insertions [Gilbert and Belliardo, 2022] and recent lateral gene transfer [Dunning et al., 2019].

## 4 Materials and Methods

### 4.1 Code structure

To identify sequences that are present in sample *P* and absent in sample *A*, GenomeDelta requires an assembly in FASTA format (*P*) and sequencing reads in FASTQ format (*A*).

GenomeDelta proceeds in several steps relying on widely used tools. First, GenomeDelta aligns the reads from *A* to *P* using bwa mem (v 0.7.17, Li and Durbin [2010]). Note that it is important to use an algorithm that performs a local alignment of the reads (such as bwa mem), otherwise the boundaries of the coverage gaps may be inferred less accurately. The mapped reads are sorted with samtools (v 1.17, [Li et al., 2009]), converted to bam files, and the coverage is computed with samtools depth. Next coverage gaps, i.e. regions with zero coverage (default threshold), are identified. To allow for some spurious mapping of reads, GenomeDelta enables merging adjacent coverage gaps separated by a maximum distance of *d* (fig. S5). All coverage gaps having a minimum size (per default 1000 bp) are extracted into a fasta-file with bedtools and a bias score is computed for each gap. The score is computed as 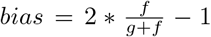 where *f* is the coverage in the 10kb regions flanking the gap (on both sides) and *g* the average coverage of the genome (fig. S1). Bedtools is used to compute the coverage in the regions flanking the gaps (v 2.30.0, [Quinlan and Hall, 2010]). The bias score ranges from −1 to 1, with 0 indicating an unbiased coverage in the flanking regions (i.e. flanking regions have the same coverage as the genomic average). A sample-specific repetitive sequence will lead to several coverage gaps, where each gap correspond to one insertion of the repetitive sequence (e.g. the dispersed insertions of a TE family). GenomeDelta derives a single consensus sequence for each repetitive sequence. To do this, GenomeDelta, extracts the sequence of each coverage gap, clusters them based on a similarity search with BLAST (v 2.6.0, [Chaisson and Tesler, 2012]) and a Python script (blast2clusters.py). We use a minimum of 3 sequences per cluster. For each cluster, a multiple sequence alignment (MSA) is generated with MUSCLE (v 3.8.1551, [Edgar, 2004]) and the consensus sequence is derived using a Python script “MSA2consensus.py”. The coverage bias of each cluster is calculated as the median of the individual biases of the clustered sequences.

As main output GenomeDelta provides two fasta files: 1) GD-candidates.fasta contains the consensus sequences of the repetitive clusters and 2) GD-non-repetitive.fasta has the sequences of the non-repetitive coverage gaps. Furthermore, a bed file is generated, which includes the genomic coordinates and the coverage bias of each coverage gap. Additionally, GenomeDelta allows to access all generated intermediate files in the output folder. Finally, GenomeDelta provides a plot summarizing the properties of the repetitive clusters, i.e the number of sequences in the cluster (e.g. the copy numbers of a TE family), the average length of the sequences and the average coverage bias. The plot is computed with the R package ggplot2 (v 3.4.4, [Wickham, 2016]).

### 4.2 Simulated data

To simulate artificial data, we used the chromosome arm 2R of *D. melanogaster* from the assembly GCA000001215.4 (strain ISO1 [Hoskins et al., 2015]) as reference sequence (sample *A*). We randomly inserted repetitive sequences into this reference sequences with SimulaTE (v 1.31, [Kofler, 2018]). Artificial reads were generated with a Python script (create-reads.py). Artificial reads that capture properties of ancient DNA were simulated with Gargammel (v3, [Renaud et al., 2017]) using 10% bacterial contamination (*Wolbachia pipientis*, an endosymbiont of *D. melanogaster*), 8% contamination with modern *D. melanogaster* DNA and 82% DNA of interest. The similarity between the observed consensus sequence and the simulated one was computed with BLAST 2.6.0 [Camacho et al., 2009].

### 4.3 Validation with real data

To validate GenomeDelta with real data we used short-reads of the strain H10 (1815, Sweden) as well as an assembly of strain Pi2 [Shpak et al., 2023, Wierzbicki et al., 2022]. We randomly subsampled the reads with Rasusa v0.8.0 [Hall, 2022] to obtain different coverages.

### 4.4 Z. tritici

We used DeviaTE (v0.3.8) [Weilguny and Kofler, 2019] to estimate the copy numbers of different TEs, including Rosetta (the novel TE identified by GenomeDelta), in *Z. tritici* strains collected from diverse geographic regions. Short reads were aligned to the sequences of the TEs and to three single copy genes (*MYCGRDRAFT-39655, MYCGRDRAFT-70396* and *MYCGRDRAFT-99758*) with bwa bwasw (version 0.7.17-r1188) [Li and Durbin, 2010]. DeviaTE estimates the copy number of a TE by normalizing the coverage of the TE by the coverage of the single copy genes [Weilguny and Kofler, 2019].

## Acknowledgments

We thank Sarah Saadain and Almoró Scarpa for testing GenomeDelta and the members of the Institute of Population Genetics for their feedback.

## Author contributions

R.P. and R.K. conceived the project. R.P. implemented the code and wrote the manuscript. A.H. validated the code. R.K. and A.H. contributed to writing the manuscript.

## Code Availability

GenomeDelta is open source and freely available at https://github.com/rpianezza/GenomeDelta. A de-tailed manual with installation instructions (https://github.com/rpianezza/GenomeDelta?tab=readme-ov-file#readme) as well as a walkthrough (https://github.com/rpianezza/GenomeDelta/blob/main/walkthrough/walkthrough.md) are available. Code for the validation can be found on GitHub.

## References

I. Anaconda. Conda: A cross-platform, language-agnostic binary package manager, 2012. URL https://github.com/conda/conda. Version 4.10.3.

D. Anxolabéhère, M. G. Kidwell, and G. Periquet. Molecular characteristics of diverse populations are consistent with the hypothesis of a recent invasion of Drosophila melanogaster by mobile P elements. Molecular Biology and Evolution, 5(3):252–269, 1988.

T. Badet, U. Oggenfuss, L. Abraham, B. A. McDonald, and D. Croll. A 19-isolate reference-quality global pangenome for the fungal wheat pathogen zymoseptoria tritici. BMC Biol, 18(12), 2020.

P. M. Benham and R. C. Bowie. Natural history collections as a resource for conservation genomics: Understanding the past to preserve the future. Journal of Heredity, 114(4):367–384, 2023.

A. Bergström, D. W. Stanton, U. H. Taron, L. Frantz, M.-H. S. Sinding, E. Ersmark, S. Pfrengle, M. Cassatt-Johnstone, O. Lebrasseur, L. Girdland-Flink, et al. Grey wolf genomic history reveals a dual ancestry of dogs. Nature, 607(7918):313–320, 2022.

E. Bonnivard, C. Bazin, B. Denis, and D. Higuet. A scenario for the hobo transposable element invasion, deduced from the structure of natural populations of Drosophila melanogaster using tandem TPE repeats. Genetical Research, 75(1):13–23, 2000.

A. Bucheton, C. Vaury, M. C. Chaboissier, P. Abad, A. Pélisson, and M. Simonelig. I elements and the Drosophila genome. Genetica, 86(1-3):175–90, 1992.

C. Camacho, G. Coulouris, V. Avagyan, N. Ma, J. Papadopoulos, K. Bealer, and T. L. Madden. Blast+: architecture and applications. BMC bioinformatics, 10:1–9, 2009.

S. E. Celniker and G. M. Rubin. The drosophila melanogaster genome. Annual review of genomics and human genetics, 4(1):89–117, 2003.

M. J. Chaisson and G. Tesler. Mapping single molecule sequencing reads using basic local alignment with successive refinement (blasr): application and theory. BMC bioinformatics, 13(1):238, 2012.

M. Chakraborty, C. Chang, D. Khost, J. A. J Vedanayagam, Y. Liao, K. Montooth, C. Meiklejohn, A. Larracuente, and J. Emerson. Evolution of genome structure in the Drosophila simulans species complex. Genome Research, 31:380–396, 2021.

B. Chénais, A. Caruso, S. Hiard, and N. Casse. The impact of transposable elements on eukaryotic genomes: from genome size increase to genetic adaptation to stressful environments. Gene, 509(1):7–15, 2012.

S. B. Daniels, A. Chovnick, and I. A. Boussyy. Distribution of hobo transposable elements in the genus Drosophila. Molecular Biology and Evolution, 7(6):589–606, 1990.

L. T. Dunning, J. K. Olofsson, C. Parisod, R. R. Choudhury, J. J. Moreno-Villena, Y. Yang, J. Dionora, W. P. Quick, M. Park, J. L. Bennetzen, et al. Lateral transfers of large dna fragments spread functional genes among grasses. Proceedings of the National Academy of Sciences, 116(10):4416–4425, 2019.

R. C. Edgar. Muscle: multiple sequence alignment with high accuracy and high throughput. Nucleic acids research, 32(5):1792–1797, 2004.

D. G. Eickbush and T. H. Eickbush. Vertical transmission of the retrotransposable elements R1 and R2 during the evolution of the Drosophila melanogaster species subgroup. Genetics, 139(2):671–684, 1995.

S. F. Elena, L. Ekunwe, N. Hajela, S. A. Oden, and R. E. Lenski. Distribution of fitness effects caused by random insertion mutations in Escherichia coli. Genetica, 102-103:349–358, 1998.

A. Feurtey, C. Lorrain, M. C. McDonal, A. Milgate, P. S. Solomon, R. Warren, G. Puccetti, G. Scalliet, S. F. F. Torriani, L. Gout, T. C. Marcel, F. Suffert, J. Alassimone, A. Lipzen, Y. Yoshinaga, C. Daum, K. Barry, I. V. Grigoriev, S. B. Goodwin, A. Genissel, M. F. Seidl, E. H. Stukenbrock, M.-H. Lebrun, G. H. J. Kema, B. A. McDonald, and D. Croll. A thousand-genome panel retraces the global spread and adaptation of a major fungal crop pathogen. Nature Communications, 14(1059), 2023a.

A. Feurtey, C. Lorrain, M. C. McDonald, A. Milgate, P. S. Solomon, R. Warren, G. Puccetti, G. Scalliet, S. F. Torriani, L. Gout, et al. A thousand-genome panel retraces the global spread and adaptation of a major fungal crop pathogen. Nature communications, 14(1):1059, 2023b.

C. Gilbert and C. Belliardo. The diversity of endogenous viral elements in insects. Current Opinion in Insect Science, 49:48–55, 2022.

C. Goubert, R. J. Craig, A. F. Bilat, V. Peona, A. A. Vogan, and A. V. Protasio. A beginner’s guide to manual curation of transposable elements. Mobile DNA, 13(1):7, 2022.

M. B. Hall. Rasusa: Randomly subsample sequencing reads to a specified coverage. Journal of Open Source Software, 7(69):3941, 2022. doi: 10.21105/joss.03941. URL https://doi.org/10.21105/joss.03941.

G. Hernández-Alonso, J. Ramos-Madrigal, H. van Grouw, M. M. Ciucani, E. L. Cavill, M.-H. S. Sinding, S. Gopalakrishnan, G. Pacheco, and M. T. P. Gilbert. Redefining the evolutionary history of the rock dove, columba livia, using whole genome sequences. Molecular Biology and Evolution, 40(11):msad243, 2023.

D. R. Hoen, G. Hickey, G. Bourque, J. Casacuberta, R. Cordaux, C. Feschotte, A.-S. Fiston-Lavier, A. Hua-Van, R. Hubley, A. Kapusta, E. Lerat, F. Maumus, D. D. Pollock, H. Quesneville, A. Smit, T. J. Wheeler, T. E. Bureau, and M. Blanchette. A call for benchmarking transposable element annotation methods. Mobile DNA, 6(1), Aug. 2015. ISSN 1759-8753. doi: 10.1186/s13100-015-0044-6. URL http://dx.doi.org/10.1186/s13100-015-0044-6.

C. J. Hogg, L. Silver, E. A. McLennan, and K. Belov. Koala genome survey: An open data resource to improve conservation planning. Genes, 14(3):546, 2023.

R. A. Hoskins, J. W. Carlson, K. H. Wan, S. Park, I. Mendez, S. E. Galle, B. W. Booth, B. D. Pfeiffer, R. A. George, R. Svirskas, et al. The release 6 reference sequence of the drosophila melanogaster genome. Genome research, 25(3):445–458, 2015.

R. N. Johnson, D. O’Meally, Z. Chen, G. J. Etherington, S. Y. Ho, W. J. Nash, C. E. Grueber, Y. Cheng, C. M. Whittington, S. Dennison, et al. Adaptation and conservation insights from the koala genome. Nature genetics, 50(8):1102–1111, 2018.

V. V. Kapitonov and J. Jurka. Molecular paleontology of transposable elements in the Drosophila melanogaster genome. Proceedings of the National Academy of Sciences of the United States of America, pages 6569–74, 2003. ISSN 0027-8424.

M. G. Kidwell. Evolution of hybrid dysgenesis determinants in Drosophila melanogaster. Proceedings of the National Academy of Sciences, 80(6):1655–1659, 1983.

R. Kofler. Simulate: simulating complex landscapes of transposable elements of populations. Bioinformatics, 34(8):1419–1420, 2018.

R. Kofler, T. Hill, V. Nolte, A. Betancourt, and C. Schlötterer. The recent invasion of natural Drosophila simulans populations by the P-element. PNAS, 112(21):6659–6663, 2015.

E. V. Koonin, K. S. Makarova, and L. Aravind. Horizontal gene transfer in prokaryotes: quantification and classification. Annual Reviews in Microbiology, 55(1):709–742, 2001.

P. Korlević, E. McAlister, M. Mayho, A. Makunin, P. Flicek, and M. K. Lawniczak. A minimally morphologically destructive approach for dna retrieval and whole-genome shotgun sequencing of pinned historic dipteran vector species. Genome biology and evolution, 13(10):evab226, 2021.

H. Li and R. Durbin. Fast and accurate long-read alignment with Burrows-Wheeler transform. Bioinformatics, 26(5):589–595, 2010.

H. Li, B. Handsaker, A. Wysoker, T. Fennell, J. Ruan, N. Homer, G. Marth, G. Abecasis, R. Durbin, and 1000 Genome Project Data Processing Subgroup. The Sequence Alignment/Map format and SAMtools. Bioinformatics, 25(16):2078–2079, Aug. 2009.

J. Nelson, A. Slicko, and Y. Yamashita. The retrotransposon R2 maintains Drosophila ribosomal DNA repeats. PNAS, 120(23):e2221613120, 2023.

L. Orlando, R. Allaby, P. Skoglund, C. Der Sarkissian, P. W. Stockhammer, M.C. Ávila Arcos, Q. Fu, J. Krause, E. Willerslev, A. C. Stone, and C. Warinner. Ancient dna analysis. Nature Reviews Methods Primers, 2021. doi: 10.1038/s43586-020-00011-0. URL http://dx.doi.org/10.1038/s43586-020-00011-0.

M. Parejo, D. Wragg, D. Henriques, J.-D. Charrière, and A. Estonba. Digging into the genomic past of swiss honey bees by whole-genome sequencing museum specimens. Genome Biology and Evolution, 12(12):2535–2551, 2020.

E. Pasyukova, S. Nuzhdin, T. Morozova, and T. Mackay. Accumulation of transposable elements in the genome of Drosophila melanogaster is associated with a decrease in fitness. Journal of Heredity, 95(4):284–290, 2004.

J. Peccoud, V. Loiseau Cordaux, and C. Gilbert. Massive horizontal transfer of transposable elements in insects. Proc Natl Acad Sci U S A, 114(18):4721–26, 2017.

J. Peccoud, R. Cordaux, and C. Gilbert. Analyzing horizontal transfer of transposable elements on a large scale: challenges and prospects. BioEssays, 40(2):1700177, 2018.

R. Pianezza, A. Scarpa, P. Narayanan, S. Signor, and R. Kofler. Spoink, a ltr retrotransposon, invaded d. melanogaster populations in the 1990s. bioRxiv, Nov. 2023. doi: 10.1101/2023.10.30.564725. URL http://dx.doi.org/10.1101/2023.10.30.564725.

R. Pianezza, A. Scarpa, A. Haider, S. Signor, and R. Kofler. Unveiling the complete invasion history of d. melanogaster: three horizontal transfers of transposable elements in the last 30 years. bioRxiv, 2024. URL https://www.biorxiv.org/content/early/2024/04/28/2024.04.25.591091.

H. Quesneville, C. M. Bergman, O. Andrieu, D. Autard, D. Nouaud, M. Ashburner, and D. Anxolabehere. Combined evidence annotation of transposable elements in genome sequences. PLoS Comp. Biol., 1(2):166–175, 2005.

A. R. Quinlan and I. M. Hall. BEDTools: a flexible suite of utilities for comparing genomic features. Bioinformatics (Oxford, England), 26(6):841–842, 2010.

C. J. Raxworthy and B. T. Smith. Mining museums for historical dna: advances and challenges in museomics. Trends in Ecology and Evolution, 2021. doi: 10.1016/j.tree.2021.07.009. URL http://dx.doi.org/10.1016/j.tree.2021.07.009.

G. Renaud, K. Hanghøj, E. Willerslev, and L. Orlando. gargammel: a sequence simulator for ancient dna. Bioinformatics, 33(4):577–579, 2017.

F. Rodriguez and I. R. Arkhipova. An Overview of Best Practices for Transposable Element Identification, Classification, and Annotation in Eukaryotic Genomes, page 1–23. Springer US, Dec. 2022. ISBN 9781071628836. doi: 10.1007/978-1-0716-2883-61. URL http://dx.doi.org/10.1007/978-1-0716-2883-6_1.

P. Sarkies, M. E. Selkirk, J. T. Jones, V. Blok, T. Boothby, B. Goldstein, B. Hanelt, A. Ardila-Garcia, N. M. Fast, P. M. Schiffer, C. Kraus, M. J. Taylor, G. Koutsovoulos, M. L. Blaxter, and E. A. Miska. Ancient and novel small RNA pathways compensate for the loss of piRNAs in multiple independent nematode lineages. PLoS Biol., 13(2):1–20, 2015.

A. Scarpa, R. Pianezza, F. Wierzbicki, and R. Kofler. Genomes of historical specimens reveal multiple invasions of ltr retrotransposons in drosophila melanogaster populations during the 19th century. bioRxiv, 2023. doi: 10.1101/2023.06.06.543830.

F. Schwarz, F. Wierzbicki, K.-A. Senti, and R. Kofler. Tirant Stealthily Invaded Natural Drosophila melanogaster Populations during the Last Century. Molecular Biology and Evolution, 38(4):1482–1497, 2021.

M. Shpak, H. R. Ghanavi, J. D. Lange, J. E. Pool, and M. C. Stensmyr. Genomes from historical drosophila melanogaster specimens illuminate adaptive and demographic changes across more than 200 years of evolution. PLoS Biology, 21(10):e3002333, 2023.

S. Signor, J. Vedanayagam, B. Kim, F. Wierzbicki, R. Kofler, and E. Lai. Rapid evolutionary diversification of the flamenco locus across simulans clade drosophila species. PLoS Genet, 19, 2023.

E. H. Stukenbrock, F. G. Jørgensen, M. Zala, T. T. Hansen, B. A. McDonald, and M. H. Schierup. Wholegenome and chromosome evolution associated with host adaptation and speciation of the wheat pathogen mycosphaerella graminicola. PLoS genetics, 6(12):e1001189, 2010.

R. E. Tarlinton, J. Meers, and P. R. Young. Retroviral invasion of the koala genome. Nature, 2006. doi: 10.1038/nature04841.

S. Van Wyk, B. D. Wingfield, L. De Vos, N. A. Van der Merwe, and E. T. Steenkamp. Genome-wide analyses of repeat-induced point mutations in the ascomycota. Frontiers in Microbiology, 11:622368, 2021.

G. L. Wallau, M. F. Ortiz, and E. L. S. Loreto. Horizontal transposon transfer in eukarya: detection, bias, and perspectives. Genome Biology and Evolution, 4(8):801–811, 2012.

L. Weilguny and R. Kofler. DeviaTE: Assembly-free analysis and visualization of mobile genetic element composition. Molecular Ecology Resources, 19(5):1346–1354, 2019.

T. Wicker, F. Sabot, A. Hua-Van, J. L. Bennetzen, P. Capy, B. Chalhoub, A. Flavell, P. Leroy, M. Morgante, O. Panaud, et al. A unified classification system for eukaryotic transposable elements. Nature reviews genetics, 8(12):973–982, 2007.

H. Wickham. ggplot2: Elegant Graphics for Data Analysis. Springer Nature, Basel, Switzerland, 2016. ISBN 978-3-319-24277-4.

F. Wierzbicki, F. Schwarz, O. Cannalonga, and R. Kofler. Novel quality metrics allow identifying and generating high-quality assemblies of pirna clusters. Molecular Ecology Resources, 22(1):102–121, 2022.

H.-H. Zhang, J. Peccoud, M.-R.-X. Xu, X.-G. Zhang, and C. Gilbert. Horizontal transfer and evolution of transposable elements in vertebrates. Nature communications, 11(1):1362, 2020.

