## Supplementary Material for "GenomeDelta: detecting recent transposable element invasions without repeat library"

#### Supplementary figures

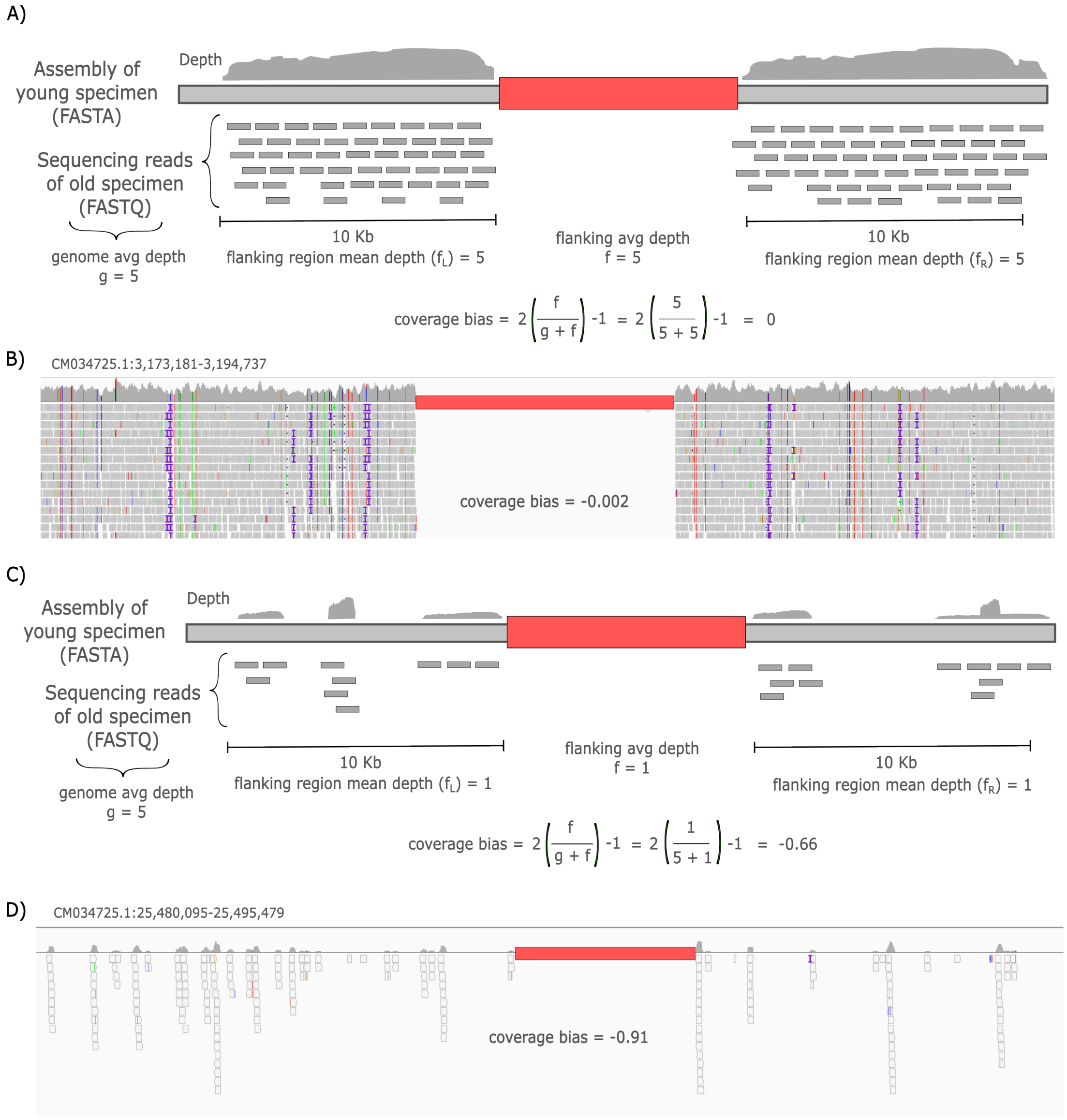


Figure 1: Overview of the coverage bias computed by GenomeDelta. Based on the coverage in the regions flanking ($f$) a sample specific sequence (red) and the average genomic coverage ($g$) the bias is computed as $bias = 2(f/(g+f)) - 1$. Illustration (A) and example (B) for an unbiased sample-specific sequence. Illustration (C) and example (D) for a biased sample-specific sequence. In these case (C,D) the coverage of the flanking regions is lower than the genomic average. Sample-specific sequences with a coverage biases $\ll 0$ or $\gg 0$ may be unreliable.

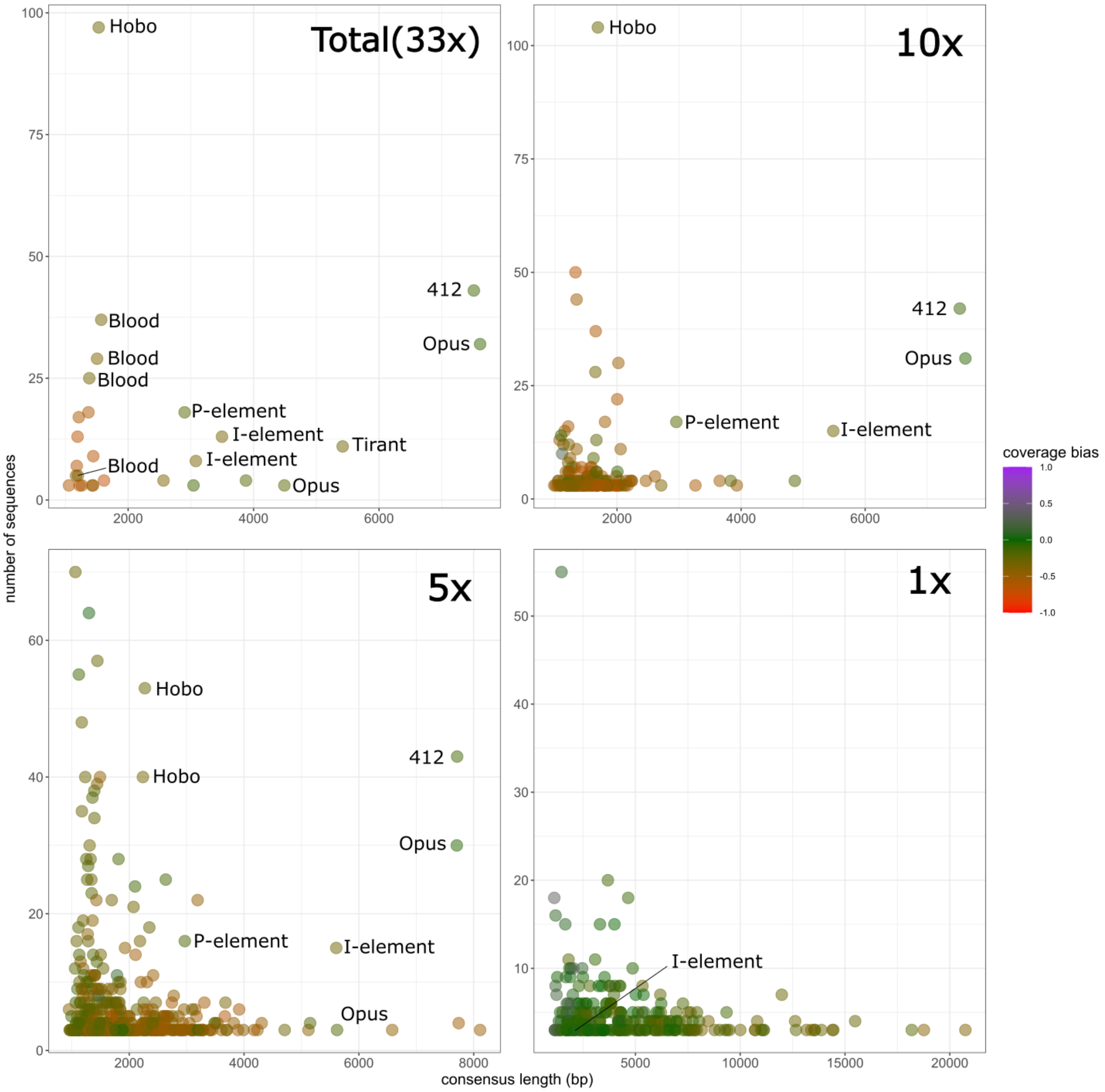


Figure 2: Effect of the coverage on the power to identify sample-specific sequences. Short-reads from historical *D. melanogaster* specimens collected in the early 1800 (H10) were aligned to an assembly of a strain collected in 1975 (Pi2) and sample-specific sequences were identified with GenomeDelta. To evaluate the performance of GenomeDelta with different coverages we subsampled the reads to coverages of 10x, 5x and 1x. With the complete data sets (33x) all 7 TEs that invaded *D. melanogaster* between 1800 and 1975 could be identified. The number of identified TE invasions decreases with the coverage. However with a coverage of 5x most invasions (5/7) were identified.

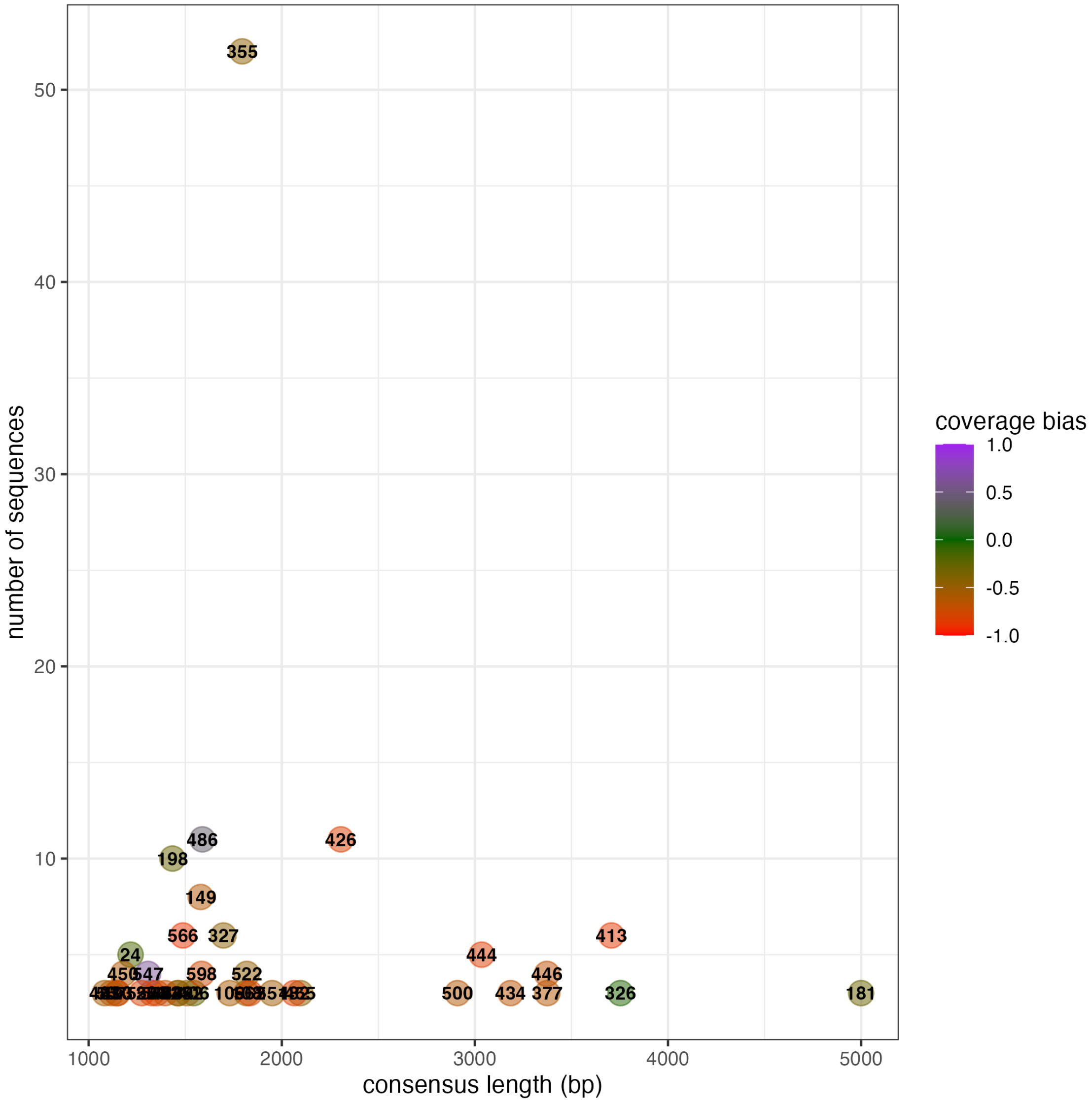


Figure 3: KoRV is not reported by GenomeDelta when two koalas having KoRV insertions are compared (short read data from a koala collected at Sunshine-Coast-M-79817 were aligned against the reference genome). None of the sample-specific sequences correspond to KoRV.

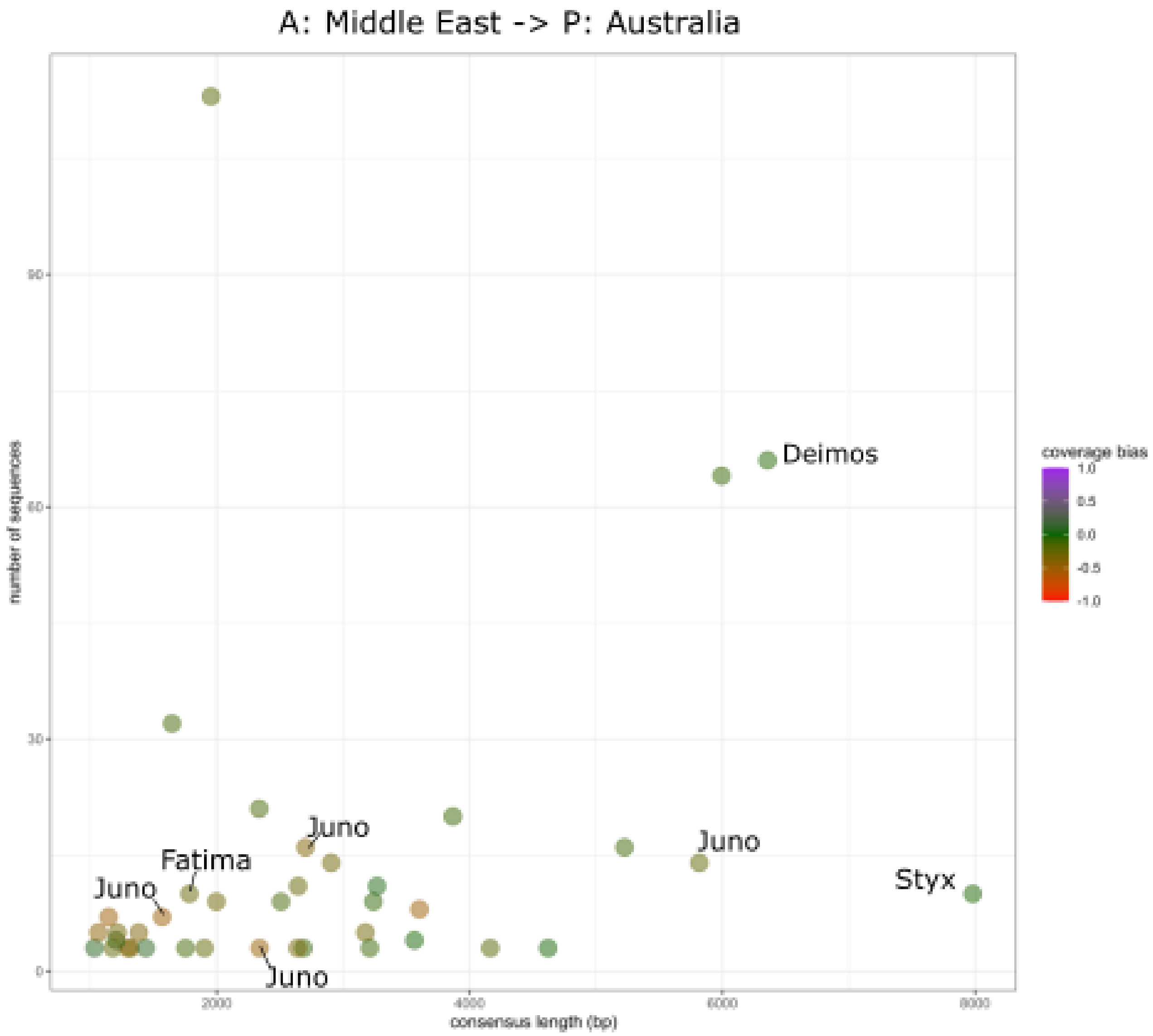


Figure 4: GenomeDelta reproduces the geographically heterogeneous distribution of several TEs (*Styx*, *Juno*, *Deimos* and *Fatima*) in *Z. tritici* reported in previous work ([Feurtey et al., 2023]). Short read data from a strain collected in Iran (SRR5194593) were aligned to an assembly of a strain collected in Australia (ERZ16268299).

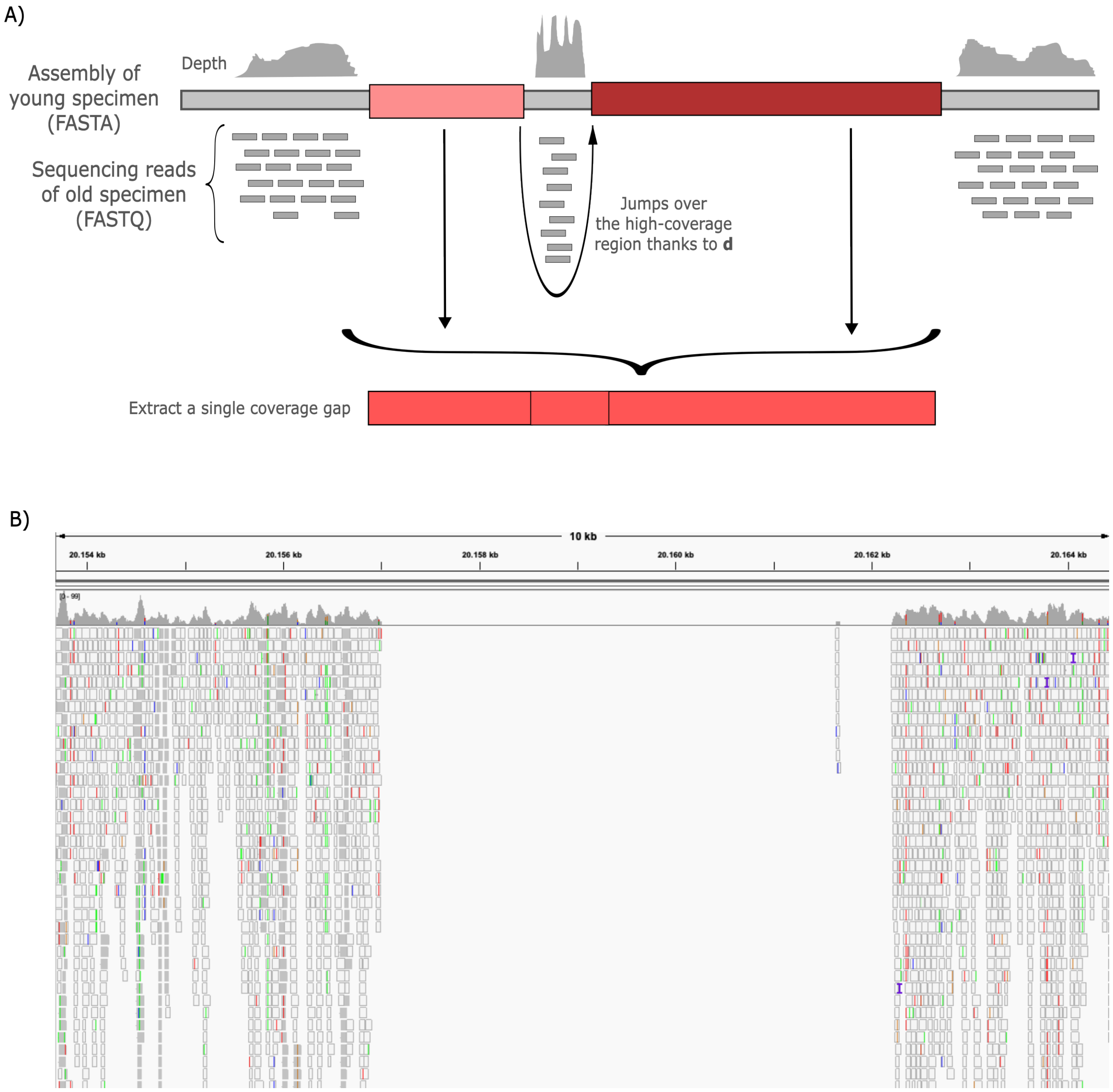


Figure 5: Overview of the parameter $d$ which specifies the maximum distance between adjacent sample-specific sequences. A) Sample specific sequences with a distance $< d$ will be merged. B) Biological example of a scenario that motivated us to introduce the parameter $d$. The coverage gap, caused by an insertion of the transposon *Spoink* in *D. melanogaster*, is interrupted by a few reads aligning within the gap (arrow). The parameter $d$ enables tolerating a few reads aligning within coverage gaps. As a result, GenomeDelta reports only one sample-specific sequence (i.e. the complete sequence of *Spoink*) instead of two using a naive approach.

#### Supplementary tables

Table 1: Validation of GenomeDelta with simulated data. We introduced 25 copies of an artificial sequence with a length of 1000bp into a template sequence and tested if the TE was accurately identified by GenomeDelta. We simulated different read coverages using either a uniform or a heterogeneous coverage (random position of reads). We also simulated properties of ancient DNA with Gargammel. We evaluated the number of true positive insertions ($TP$), the number of false positive insertions ($FP_r$, $FP_{nr}$: repetitive or non-repetitive) and the length of the reported consensus sequence [len. (bp)]. Note that many false positives are detected when the coverage is low.

| method | coverage | $TP$ | $FP_{nr}$ | $FP_r$ | len. (bp) |
|---|---|---|---|---|---|
| uniform | 1 | 25/25 | 102 | 7 | 999 |
| uniform | 5 | 25/25 | 0 | 0 | 998 |
| uniform | 10 | 25/25 | 0 | 0 | 998 |
| random | 1 | 25/25 | 4574 | 6 | 1134 |
| random | 5 | 25/25 | 0 | 0 | 999 |
| random | 10 | 25/25 | 0 | 0 | 998 |
| gargammel | 1 | 24/25 | 8873 | 40 | 1206 |
| gargammel | 5 | 25/25 | 0 | 0 | 1003 |
| gargammel | 10 | 25/25 | 0 | 0 | 998 |

Table 2: Performance of GenomeDelta with different sequencing error rates. We introduced 25 copies of an artificial sequence with a length of 5000bp into a template sequence and tested if the TE was accurately identified by GenomeDelta. We simulated different coverages with randomly distributed reads having error rates ranging from 0 to 5% (error). We evaluated the number of true positive insertions ($TP$), the number of false positive insertions ($FP_r$, $FP_{nr}$: repetitive or non-repetitive) and the length of the reported consensus sequence [len. (bp)]. Note that the error rate had little impact on the performance of GenomeDelta (apart from slightly elevated $FP_{nr}$ at low coverages).

| error | coverage | $TP$ | $FP_{nr}$ | $FP_r$ | len. (bp) |
|---|---|---|---|---|---|
| 0 | 1 | 25/25 | 607 | 2 | 5251 |
| 0 | 5 | 25/25 | 0 | 0 | 4998 |
| 0 | 10 | 25/25 | 0 | 0 | 4997 |
| 0.01 | 1 | 25/25 | 838 | 1 | 5150 |
| 0.01 | 5 | 25/25 | 0 | 0 | 4999 |
| 0.01 | 10 | 25/25 | 0 | 0 | 4999 |
| 0.05 | 1 | 25/25 | 984 | 1 | 5160 |
| 0.05 | 5 | 25/25 | 0 | 0 | 4999 |
| 0.05 | 10 | 25/25 | 0 | 0 | 4999 |

Table 3: Performance of GenomeDelta with different read lengths. We introduced 25 copies of an artificial sequence with a length of 5000bp into a template sequence and tested if the TE was accurately identified by GenomeDelta. We simulated different coverages with randomly distributed reads having lengths ranging from 50 to 200bp (length). We evaluated the number of true positive insertions ($TP$), the number of false positive insertions ($FP_r$, $FP_{nr}$: repetitive or non-repetitive) and the length of the reported consensus sequence [len. (bp)]. Note that the number of false positives ($FP_{nr}$) decreased with the read length.

| length | coverage | $TP$ | $FP_{nr}$ | $FP_r$ | len. (bp) |
|---|---|---|---|---|---|
| 50 | 1 | 25/25 | 2120 | 0 | 5752 |
| 50 | 5 | 25/25 | 0 | 0 | 4999 |
| 50 | 10 | 25/25 | 0 | 0 | 4999 |
| 100 | 1 | 25/25 | 607 | 0 | 5251 |
| 100 | 5 | 25/25 | 0 | 0 | 4998 |
| 100 | 10 | 25/25 | 0 | 0 | 4997 |
| 200 | 1 | 25/25 | 194 | 0 | 5038 |
| 200 | 5 | 25/25 | 0 | 0 | 4999 |
| 200 | 10 | 25/25 | 0 | 0 | 4999 |

Table 4: Overview of the sample-specific repetitive sequences identified by GenomeDelta when reads from a *D. melanogaster* strain collected in 1815 ($H10$) are aligned to the assembly of a strain collected in 1975 (Pi2). For each sequence, we show the ID, the coverage bias, the number of insertions, the length of the insertion and the matching TE sequence as revealed by a BLAST search. Out of the sequences with a low coverage bias ($0.2 > bias > -0.2$), solely sequence 76 is not matching a TE sequence.

| ID | coverage bias | insertions | len. (bp) | matching TE |
|---|---|---|---|---|
| 1 | -0.66 | 13 | 1194 | |
| 2 | -0.59 | 17 | 1214 | |
| 3 | -0.60 | 18 | 1368 | |
| 6 | -0.44 | 5 | 1169 | |
| 7 | -0.09 | 32 | 7611 | *Opus* |
| 8 | -0.22 | 3 | 4491 | *Opus* |
| 9 | -0.65 | 3 | 1225 | |
| 11 | -0.32 | 25 | 1381 | *Blood* |
| 12 | -0.38 | 29 | 1505 | *Blood* |
| 13 | -0.70 | 4 | 1615 | |
| 14 | -0.12 | 43 | 7511 | *412* |
| 15 | -0.35 | 13 | 3496 | *I-element* |
| 17 | -0.33 | 37 | 1571 | *Blood* |
| 19 | -0.64 | 97 | 1531 | *Hobo* |
| 28 | -0.21 | 18 | 2901 | *P-element* |
| 41 | -0.29 | 11 | 5420 | *Tirant* |
| 46 | -0.65 | 7 | 1181 | |
| 48 | -0.65 | 9 | 1444 | |
| 49 | -0.36 | 5 | 1199 | *Blood* |
| 67 | -0.35 | 8 | 3080 | *I-element* |
| 68 | -0.48 | 3 | 1442 | |
| 76 | -0.12 | 3 | 3042 | |
| 88 | -0.20 | 4 | 3878 | |
| 179 | -0.39 | 4 | 2564 | |
| 185 | -0.51 | 3 | 1262 | |
| 353 | -0.60 | 3 | 1057 | |
| 606 | -0.64 | 3 | 1426 | |

### References

A. Feurtey, C. Lorrain, M. C. McDonal, A. Milgate, P. S. Solomon, R. Warren, G. Puccetti, G. Scalliet, S. F. F. Torriani, L. Gout, T. C. Marcel, F. Suffert, J. Alassimone, A. Lipzen, Y. Yoshinaga, C. Daum, K. Barry, I. V. Grigoriev, S. B. Goodwin, A. Genissel, M. F. Seidl, E. H. Stukenbrock, M.-H. Lebrun, G. H. J. Kema, B. A. McDonald, and D. Croll. A thousand-genome panel retraces the global spread and adaptation of a major fungal crop pathogen. *Nature Communications*, 14(1059), 2023.
